# Multi-camera Simultaneous Total Internal Reflection and Interference Reflection Microscopy

**DOI:** 10.1101/2024.08.28.610099

**Authors:** Jeffrey O. Spector, Jiayi Chen, Antonina Roll-Mecak

**Affiliations:** Cell Biology and Biophysics Unit, National Institute of Neurological Disorders and Stroke, Bethesda, MD 20892, U.S.A; Biochemistry and Biophysics Center, National Heart, Lung and Blood Institute, Bethesda, MD 20892, U.S.A

**Keywords:** Interference Reflection Microscopy, Label-free imaging, Scattering

## Abstract

Interference Reflection Microscopy (IRM) is an optical technique that relies on the interference between the reflected light from an incident beam as it passes through materials of different refractive indices. This technique has been successfully used to image microtubules, biologically important biofilaments with a diameter of 25 nm. However, it is often desirable to image both the microtubule and microtubule interacting proteins simultaneously. Here we present a simple modification to a standard multi-color total internal reflection fluorescence (TIRF) microscope that enables simultaneous high-speed IRM and single molecule TIRF imaging. Our design utilizes a camera for each channel (IRM and TIRF) allowing independent optimization of camera parameters for the two different modalities. We illustrate its application by imaging unlabeled microtubules and GFP-labeled end-binding protein EB1 which forms comets on the tips of polymerizing microtubules. Our design is easily implemented, and with minimal cost, making it accessible to any laboratory with an existing fluorescence microscope.

## Introduction

Microtubules are cylindrical biopolymers with a diameter of 25 nm. They are highly dynamic and play important roles in many fundamental cellular processes, ranging from cell division to cellular motility. Owing to their biological importance and their sub-diffraction limited size they have been imaged extensively with many modalities. The first study imaging directly the dynamic nature of microtubules used darkfield imaging [1] These were followed by studies using Differential Interference Contrast (DIC) microscopy [2], which was employed in several seminal studies into the assembly kinetics of microtubules [2, 3]. The success of these two imaging techniques revealed that, although sub-diffraction limited, microtubules contain enough mass in a diffraction-limited spot to be amenable to scattering-based imaging. However, once it was possible to fluorescently label tubulin, the primary mode of imaging microtubules switched to fluorescence. For *in vitro* microtubule dynamics assays, which require µM concentrations of soluble tubulin, total internal reflection (TIRF) imaging became the primary imaging method. [4–11] TIRF imaging has the advantage of very low background and the capability to easily multiplex several colors with the use of a multi bandpass dichroic in the excitation/emission path of the microscope. However, in recent years there has been a resurgence in label-free imaging, using either interferometric scattering microscopy (iSCAT) [12] or IRM [13, 14]. IRM imaging in particular has become a popular choice because it is low cost compared to TIRF microscopy and is relatively simple to adapt to existing fluorescent microscopes. The ability to image microtubules label-free has many advantages. First, fluorescent microtubules are susceptible to photo damage [15]. Second, understanding the roles played by tubulin post-translational modifications and isoforms requires the use of engineered recombinant tubulin [16], and this tubulin is not always available in the quantities necessary for fluorescent labelling which usually results in large losses of material. Third, the fluorescent label itself can affect the polymerization properties of the tubulin. However, in many cases it is desirable to image, in addition to the microtubule, also the proteins that interact with them to understand their localization and dynamics. Most of these proteins are too small to be easily visible by IRM, and they frequently need to be added at high concentrations. There have been several attempts to combine IRM and TIRF [17, 18] each with their own advantages and drawbacks. Generally, the imaging is performed sequentially not simultaneously, or using a single camera with a splitter to image two emission channels together on the same detector. Simultaneous imaging is desirable to capture rapid events such as microtubule associated protein (MAP) binding and unbinding, but the use of a single camera does not allow optimization of imaging conditions for both TIRF and IRM. We present here a method that is simple and inexpensive to implement that combines IRM and TIRF imaging for simultaneous visualization of dynamic microtubules and microtubule associated proteins using independent detectors for each channel.

## Materials and Methods

### Simultaneous IRM and TIRF imaging Setup

Interference reflection microscopy forms an image by the interference between the light that is scattered by the sample and the light that reflects off the sample-substate interface. IRM was originally used for imaging cell adhesion to glass [19]. In the case of *in vitro* microtubule assays, the substrate is a glass coverslip, and the scattering sample is a microtubule close to the coverslip surface. The most widely used implementation of IRM for microtubule imaging was presented by Mahamdeh [13] wherein the microscope dichroic was replaced by a 50/50 beam splitter which yields excellent results for imaging only microtubules (Figure 1A). Coupling this implementation with fluorescence imaging is challenging because the 50/50 beam splitter drastically reduces the amount of signal that can be collected in the fluorescence channel. We sought to develop a method that was capable of simultaneous IRM and fluorescence imaging without the signal reduction introduced by the 50/50 beam splitter. A solution to this problem was presented by Tuna *et. al.* [20]. In that study, the 50/50 beam splitter was replaced with a 10/90 (R:T) and the fluorescence and IRM channels were imaged on different parts of the same camera. This implementation requires much higher initial laser and lamp powers because only 10% of the excitation light is being directed towards the sample. The use of a single camera to image both channels also does not allow the independent optimization of each channel (e.g. exposure time, camera gain), and one is forced to trade off the image quality in each channel to find a middle ground. Lastly, this design also requires sacrificing at least half of the achievable field of view in each channel. A second implementation forgoes the use of a beam splitter and instead uses micromirrors [18]. This design also uses a single camera to image both the fluorescence and scattering channels. IRM has been successfully combined with optical tweezers [17]; however, it was implemented using a blue LED which can photodamage microtubules over prolonged exposure. We therefore sought to implement simultaneous TIRF and IRM imaging on two separate cameras, maintaining the ability to image single molecules in the fluorescence channel and utilizing the entire camera field of view (Figure 1B). Instead of changing the standard TIRF dichroic for a beam splitter we looked for a standard multi-bandpass dichroic that would be compatible with both TIRF and IRM imaging. Using a blue LED for IRM would yield the best IRM image because shorter wavelengths lead to a stronger scattering signal, however that could cause photodamage over long periods and time, and in addition would preclude the use of the 488 nm excitation line for simultaneous imaging. We therefore used a light source that is centered at 625 nm so that we could easily image the green fluorescence channel (emission ∼ 525 nm) with minimal photodamage from the IRM light. An ideal filter spectrum is presented in Figure 2A. We searched the transmission spectra for a commercially available dichroic that possessed ∼ 50% transmission in the 600-650 nm range. Our multicolor TIRF setup uses a Di01-R405/488/561/635 from Semrock (Figure 2B). This dichroic is essentially a 50/50 beam splitter in a small range centered around 625 nm (Figure 2C). This range coincides with the peak of commercially available LED sources. In our case, we used an M617L3 LED from ThorLabs (Figure 2B,C). We also achieved good imaging with a white-light LED (Lumencor SOLA-SE-II) by adding an excitation bandpass filter centered at 625 nm (Semrock FF01-625/26). This wavelength can be used in combination with the multiband dichroic to perform simultaneous IRM and TIRF fluorescence imaging, without loss of signal in the fluorescence channel as is the case for the designs that use a 50/50 or a 10/90 beam splitter. To allow the independent setting of the exposure time and gain for each channel, the emission channels are split, and each imaged onto a separate camera. We chose to use emCCD cameras for single molecule imaging, but standard sCMOS cameras will also work if emCCD sensitivity is not necessary. To register the IRM and fluorescence channels we image a grid of diffraction-limited spots (NanoGrid, MiralomaTech) and calculate the transformation to align them using the ImageJ gridAligner plugin (Figure 3). This yields the transformation between channels over the entire field of view and is not limited to only a few beads as in Tuna *et. al.* [20]. Images are acquired simultaneously in both the IRM and the 488 channels. The emission is split using a Di02-R594 (Semrock) and further filtered with a 514/30 for the TIRF channel and a 593LP for the IRM channel. Each detection channel is equipped with an iXON 897 emCCD camera (Andor). The signal is split using an TuCAM adapter (Andor), and the timing is controlled by an AOTF that gates the lasers and starts the image acquisition. The entire system is built around a Nikon Ti-E microscope body with a Ti-TIRF attachment, and a Thorlabs M617 LED coupled to the microscope via a Nikon EPI LED attachment module. The dichroic turret contains the Di01- 405/488/561/635 dichroic (Semrock). IRM illumination intensity is manually controlled and set to yield a good IRM image (∼ 30%). The 488 laser is set to 20 mW before being coupled to the TIRF arm via an optical fiber. The entire system was run by micromanager. [21] For a complete list of parts see Table 1.

**Figure 1.**
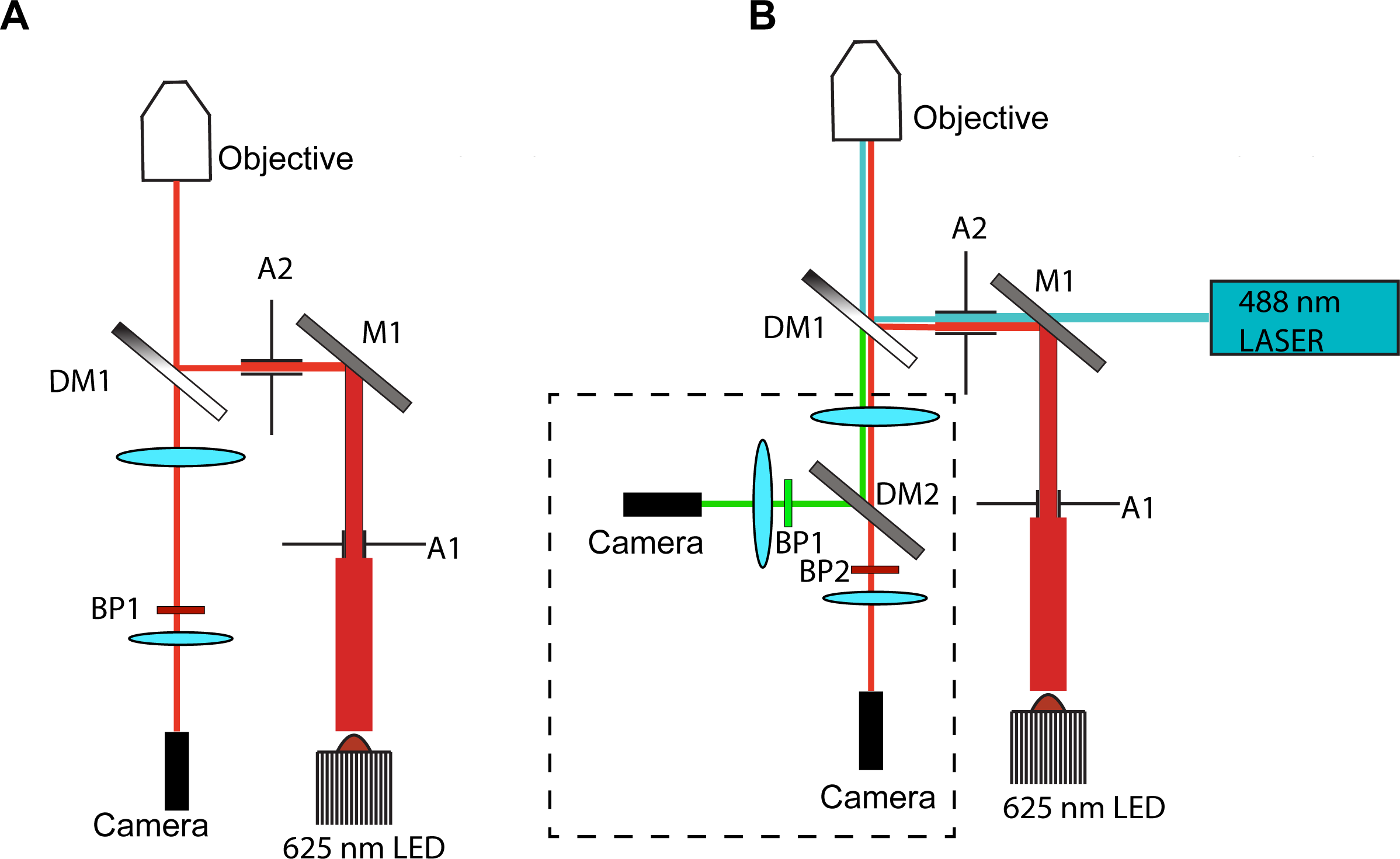
Schematics of an IRM only and IRM/Fluorescence imaging systems. A) A conventional setup for single color IRM imaging. A1, adjustable iris aperture, M1, mirror 1, A2, adjustable iris aperture, DM1, 50/50 (or 10/90) beam splitter, BP1, bandpass filter FF593/LP, camera = Andor Ixon 897 emCCD. B) Design for a simultaneous IRM/TIRF microscope. A1, adjustable iris aperture, M1, combining mirror (90:10 T:R), A2, adjustable iris aperture, DM1, quad band dichroic Di01-R405/488/561/635, DM2, Emission splitting mirror Di02-R594, BP1, bandpass filter for fluorescence channel (525/50), BP2, bandpass filter for IRM channel (593/LP). Camera, Andor Ixon 897 emCCD. This configuration also allows for multi-channel TIRF imaging if IRM is not used. The part enclosed in the dashed lines is the Andor TuCAM multi-camera adapter.

**Figure 2.**
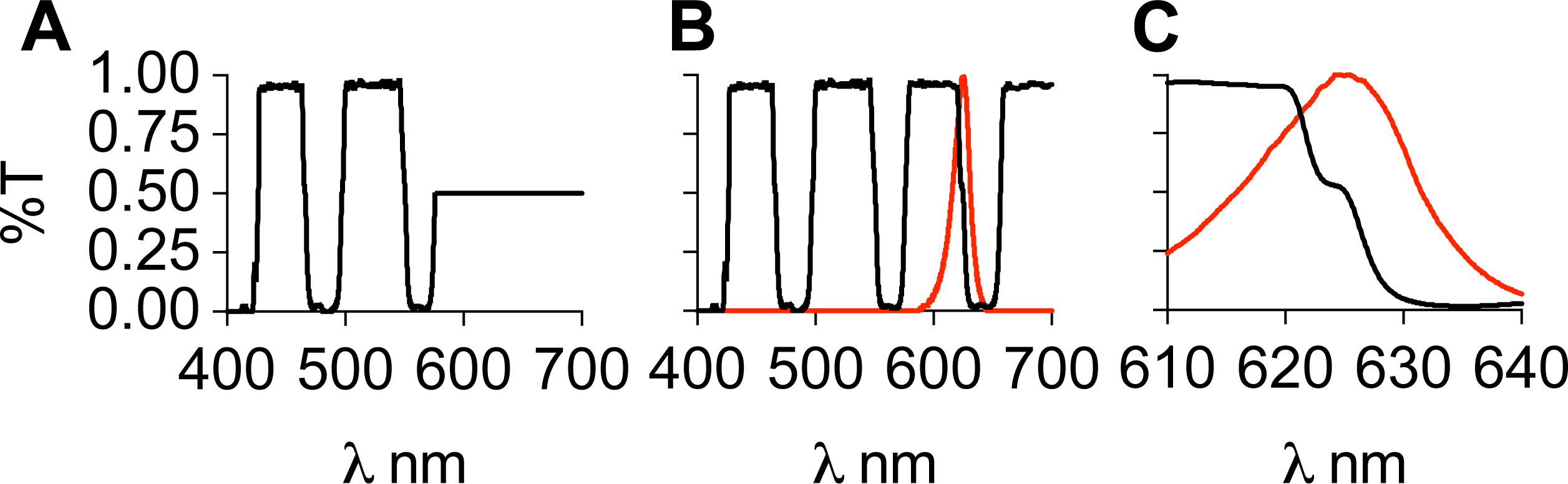
Transmission spectra of dichroic mirrors. A) The spectrum for a filter ideally suited for simultaneous IRM/TIRF imaging. The filter acts like a multi band pass dichroic up until the wavelength used for IRM imaging, at which point it acts as a 50/50 beam splitter. B) The spectrum of the Semrock Di01-R405/488/561/635 dichroic. It has the same bands as the ideal filter but also two more, centered at ∼ 600 nm and 700 nm. The red line indicates the emission spectrum of the LED used in this study. C) Zoom in between 610 nm and 640 nm. The dichroic transitions from transmitting to reflecting with a 50/50 shoulder located around 625. This coincides with the peak emission of the LED.

**Figure 3.**
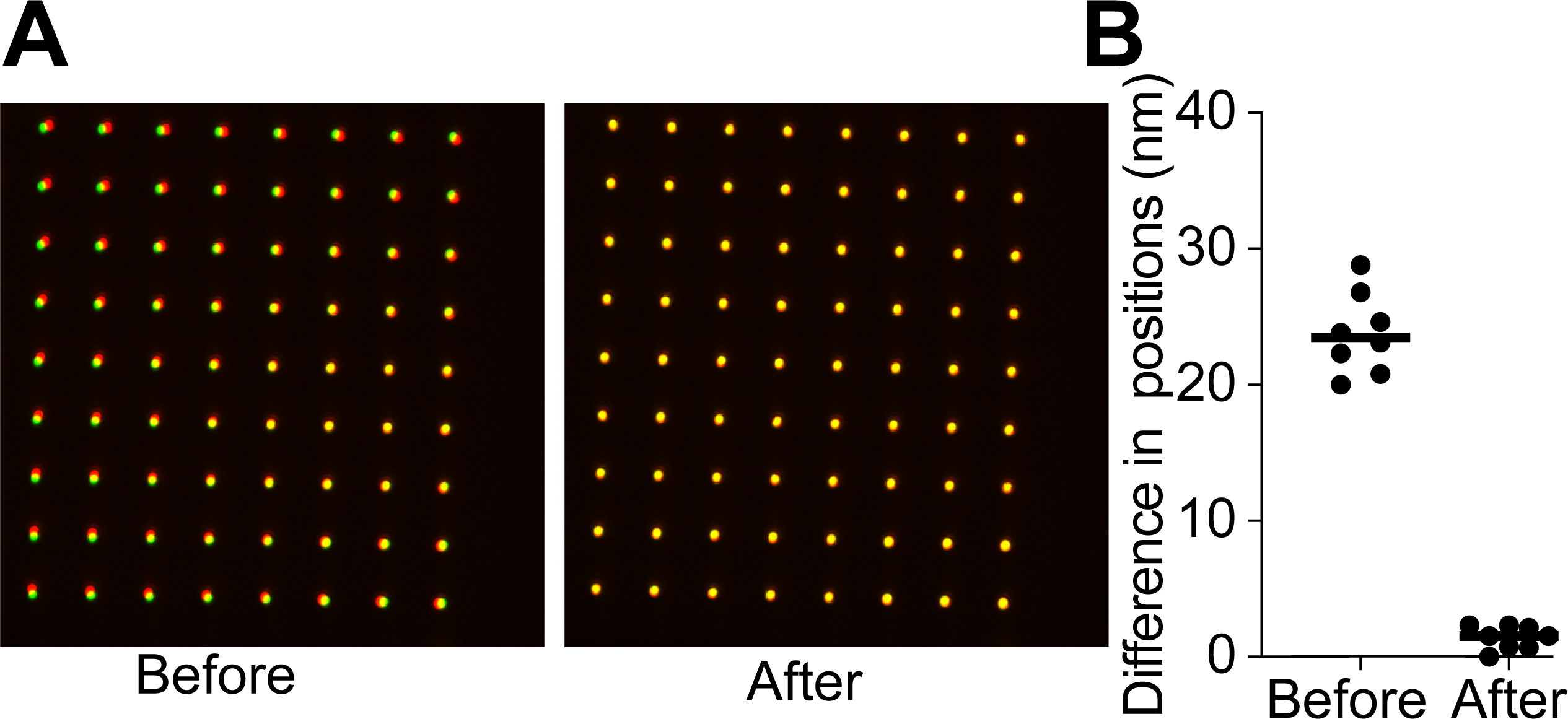
Registration between the fluoresce and IRM channels. A) An image of the NanoGrid before and after calculating the affine transformation to register the channels. Green represents the fluorescence camera and red is the IRM camera. Image is shown before and after applying the calculated transformation. B) The difference in peak positions before and after the transformation between the two channels. Before registration the difference is 25 nm After registration the alignment is accurate to 2 nm across the field of view.

**Table 1.** List of Parts.

| **Equipment** | **Manufacturer** |
| --- | --- |
| Ti-E Inverted Microscope with TIRF and LED illumination accessories | Nikon Instruments Inc. Melville,NY |
| 100X NA 1.49 TIRF objective | Nikon Instruments Inc. Melville,NY |
| LED Light source (M617L3) | ThorLabs Newton,NJ |
| Excitation Laser( 488nm Cube) | Coherent Saxonburg,PA |
| 50:50 or 10:90 Beam Splitter | ThorLabs Newton,NJ |
| IRM/Fluorescence Dichroic<br>(Di01-R405/488/561/635) | Semrock (Idex Health & Science,LLC)<br>Rochester,NY |
| Emission Filters for TuCAM<br>Di02-R594 – for splitting emission<br>FF01-593/LP – for IRM<br>FF03-525/50 – for Fluorescence | Semrock (Idex Health & Science,LLC)<br>Rochester,NY |
| Multi Camera Adapter<br>(TuCAM) | Andor USA (Oxford instruments)<br>Concord,MA |
| 2x emCCD (iXON 897) | Andor USA (Oxford instruments)<br>Concord,MA |
| NanoGrid | Miraloma Tech ,LLC San Francisco, CA |

## Microscopy Sample Preparation

Imaging chambers were prepared using a silanized coverslip attached to a slide via double sided tape [22]. Chambers were first perfused with BRB80(80 mM PIPES, pH 6.8, 1 mM EGTA, 1 mM MgCl_2_) followed by 0.1 mg/ml neutravidin for 5 minutes. Chambers were washed and GMPCPP stabilized microtubule seeds were perfused in. Seed binding to the glass was monitored by IRM until a few seeds per field of view were observable. The chamber was then washed again with BRB80, 0.1% β-mercaptoethanol, 2 mg/ml κ- casein and the final perfusion consisting of BRB80 supplemented with 50 mM KCl, 0.1% methylcellulose 4000 mPa.S, 0.5% pluronic F-127, 100 μg/ml κ-casein, 1 mM β-mercaptoethanol, 300 nM EB1-GFP and oxygen scavengers was perfused into the chamber. The chamber was maintained at 30°C with an objective heater (Bioptechs). An exposure time of 100 ms was used and images were acquired at 3 frame/second for 30 minutes.

## Results

Simultaneous IRM and TIRF were used to image dynamic microtubules in the presence of GFP-EB1. EB1 tracks the tips of growing dynamic microtubules, resulting in a “comet”- like appearance [5]. First, a background area is collected by recording a movie while rapidly moving the stage until 100 images have been acquired. The median of this stack is calculated and subtracted from the original IRM channel image to yield the background-corrected IRM image. The background corrected images were then processed through a Bandpass fouler Filter in ImageJ (Figure 4A). Figure 4B displays examples of typical kymographs of dynamic microtubule in the presence of GFP-EB1 which clearly show the size and shape of the EB1 comet in the TIRF channel and the profile of the dynamic microtubule in the IRM channel (Figure 4B,Supplmental Movie 1). Line scans in the fluorescent channel show the GFP-EB1 comets at the growing tips and illustrate the excellent signal to noise achievable while simultaneously imaging the unlabeled microtubules with IRM.

**Figure 4.**
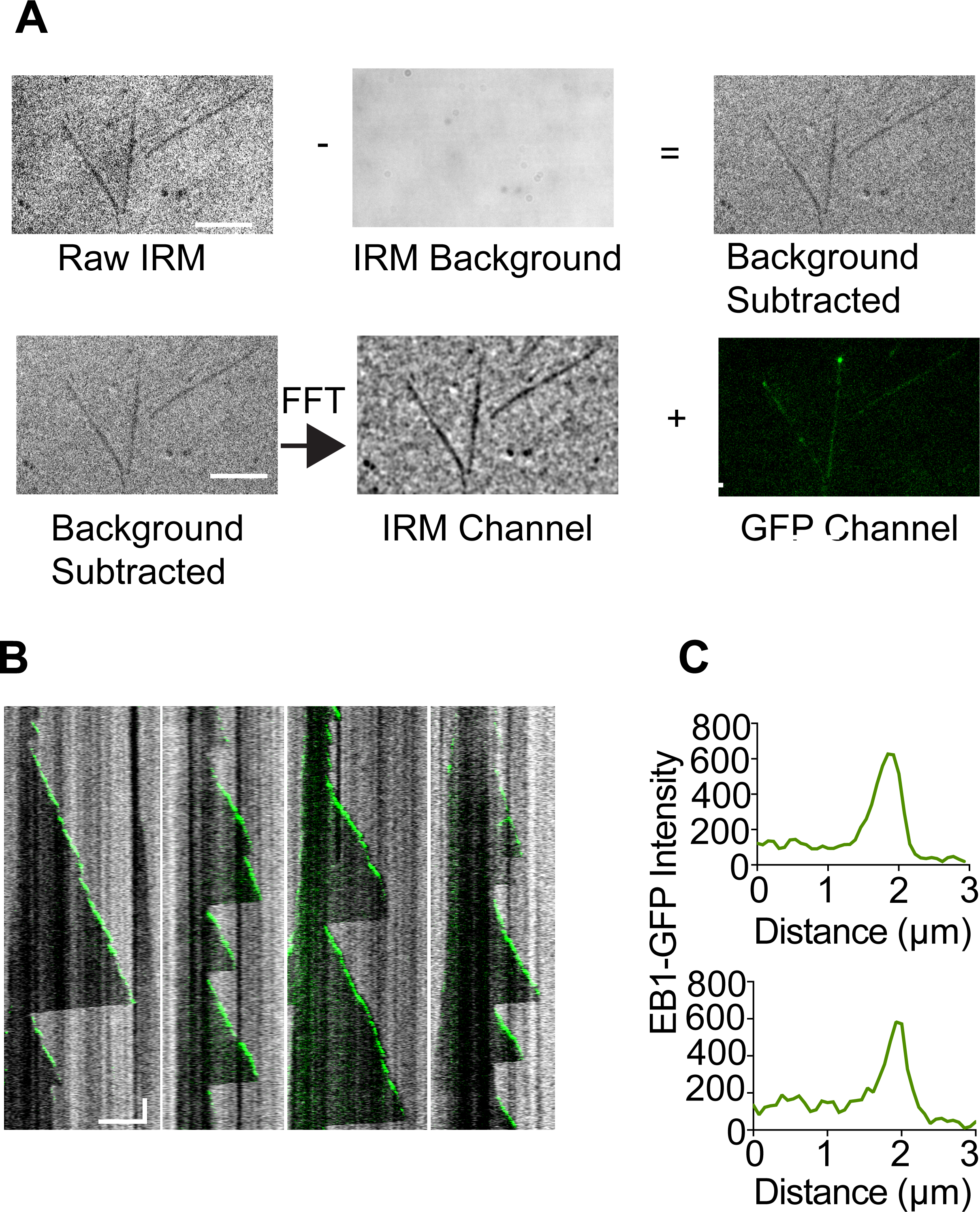
Example data acquired of dynamic microtubules imaged using simultaneous IRM and fluorescence imaging. A) The median projection of 100 frames acquired by rapidly moving the stage is subtracted from the raw IRM image. This background subtracted image is then processed in a bandpass FFT filter to yield the final image. These two steps are not strictly necessary but greatly enhance the contrast in the final image. This image is then overlayed with the fluorescence image and aligned using the transformation calculated from the NanoGrid. B) Example kymographs showing the IRM channel in grey and the GFP-EB channel in green. The dynamics (polymerization and depolymerization) of the microtubule are readily observable simultaneously with the GFP-EB1 comets. Horizontal scale bar is 5 µm and vertical scale bar represent 60 s. C) Examples of line scans across the kymographs acquired in the fluorescence channel. The canonical “comet” shape is clearly visible, and the fluorescence channel has excellent signal-to-noise.

## DISCUSSION

Here we present a simple modification to a commercial multicolor TIRF setup to incorporate simultaneous label-free IRM imaging and show its use in performing microtubule dynamics assays in the presence of a fluorescently-labeled microtubule associated protein. Imaging microtubules label-free offers several advantages. First, there is no photobleaching, typically seen when doing microtubule dynamic assays over periods of 10-15 minutes. Second, the assays do not necessitate labeling of tubulin, and are not complicated by the effects of the fluorophores on microtubule dynamics parameters. Third, the simultaneous IRM and TIRF imaging allows the observation at high temporal resolution of both microtubules and interacting partners. The method we present uses an independent camera for the IRM and TIRF channels. This has several advantages over single camera designs: it does not sacrifice field of view and allows setting the gain on each emCCD independently. This is important because the IRM signal is usually very strong and can easily saturate an emCCD camera, whereas signal in the fluorescence channel can be very dim and requires the use of gain on the emCCD camera. Because we use a LED at 625 nm, fluorophores that emit in this range are excluded; however, the use of fluorophores that emit in the far red (>650 nm) is still available. Therefore, while we demonstrated the performance of this system with GFP, it is equally compatible with fluorophores that absorb in the ∼647 nm range. Additionally, the simple addition of a third camera would allow for two-color fluorescence simultaneously with IRM imaging. An additional advantage to our design is that we can run the cameras at separate frame rates. For example, one can image microtubule dynamics at a high frame rate, and the fluorescence channel at a lower frame rate. This reduces photobleaching in the fluorescence channel and also allows imaging processes with different kinetics. In this study we applied background subtraction and a bandpass FFT filter to enhance the IRM images. We note that even without these processing steps the microtubules are visible in the raw images. (Figure 4A) If the IRM images are recorded at a high enough framerate, temporal averaging can also be used to increase the signal-to-noise [13]. Thus, this simple modification to a standard two-color TIRF microscope enables simultaneous label-free and TIRF imaging with image quality comparable to that obtained with two-color fluorescence imaging.

## CONCLUSION

Label-free imaging offers several advantages over fluorescence imaging: it virtually eliminates photobleaching; and it allows imaging of microtubules whose dynamic behavior is not impacted by the fluorescence labeling and damage due to the fluorophore photodamage. IRM microscopy is quickly becoming adapted as an easy to implement, low-cost method for imaging microtubules label free [14, 23–28]. Despite this, there is still a need to perform simultaneous IRM and fluorescence (TIRF) imaging to study the interaction between microtubules and their interacting partners. We present here a simple method using a multichannel dichroic and a LED matched to the spectrum of the dichroic in a region where the dichroic acts as a 50/05 beam splitter. This simple modification, paired with the use of dual cameras enables IRM and TIRF imaging without sacrificing field of view (as in systems that split the camera chip) and simultaneously (whereas other implementations require switching filters which decreases temporal resolution). This system has applications beyond microtubule assays and will work for any assay that requires simultaneous imaging in IRM and TIRF. By using a multi-channel dichroic, our design can easily be extended to two different fluorescent channels or can simply function as a simultaneous two-color TIRF microscope.

## Acknowledgments

J.S. thanks J. Cleary (National Institute of Neurological Disorders and Stroke) for discussion about IRM imaging. The authors would like to thank A. Szyk (National Institute of Neurological Disorders and Stroke) for preparation of the GFP-EB1. A.R-M. is supported by the intramural programs of the National Institute of Neurological Disorders and Stroke (NINDS) and the National Heart, Lung and Blood Institute (NHLBI).

## Supplemental Movie 1

A movie showing the IRM channel (left) and GFP-EB1 channel (right) acquired simultaneously. The IRM channel was background subtracted and processed by a bandpass FFT. The movie was recorded at 1 frame/3 seconds and played back at 30 fps. Individual polymerization and depolymerization events are visible in the IRM channel. The GFP channel displays the EB1 comets.

## References

1. Horio, T. and H. Hotani, Visualization of the dynamic instability of individual microtubules by dark-field microscopy. Nature, 1986. 321(6070): p. 605-607.

2. Walker, R., et al., Dynamic instability of individual microtubules analyzed by video light microscopy: rate constants and transition frequencies. The Journal of cell biology, 1988. 107(4): p. 1437–1448.

3. KristoLerson, D., T. Mitchison, and M. Kirschner, Direct observation of steady-state microtubule dynamics. The Journal of cell biology, 1986. 102(3): p. 1007–1019.

4. Gell, C., et al., Microtubule dynamics reconstituted in vitro and imaged by single-molecule fluorescence microscopy. Methods in cell biology, 2010. 95: p. 221–245.

5. Bieling, P., et al., Reconstitution of a microtubule plus-end tracking system in vitro. Nature, 2007. 450(7172): p. 1100-1105.

6. Vemu, A., et al., Severing enzymes amplify microtubule arrays through lattice GTP-tubulin incorporation. Science, 2018. 361(6404): p. eaau1504.

7. Wieczorek, M., et al., Microtubule-associated proteins control the kinetics of microtubule nucleation. Nature cell biology, 2015. 17(7): p. 907–916.

8. Komarova, Y., et al., Mammalian end binding proteins control persistent microtubule growth. Journal of Cell Biology, 2009. 184(5): p. 691–706.

9. Zanic, M., et al., EB1 recognizes the nucleotide state of tubulin in the microtubule lattice. PloS one, 2009. 4(10): p. e7585.

10. Consolati, T., et al., *Real-time imaging of single γTuRC-mediated microtubule nucleation events in vitro by TIRF microscopy*, in *Microtubules: Methods and Protocols*. 2022, Springer. p. 315-336.

11. Brouhard, G.J., et al., XMAP215 is a processive microtubule polymerase. Cell, 2008. 132(1): p. 79–88.

12. Andrecka, J., et al., Label-free imaging of microtubules with sub-nm precision using interferometric scattering microscopy. Biophysical journal, 2016. 110(1): p. 214–217.

13. Mahamdeh, M., et al., Label Free High Speed Wide Field Imaging of Single Microtubules using Interference Reflection Microscopy. Biophysical Journal, 2018. 114(3): p. 504a–505a.

14. Chen, J., et al., α-tubulin tail modifications regulate microtubule stability through selective eVector recruitment, not changes in intrinsic polymer dynamics. Developmental cell, 2021. 56(14): p. 2016–2028. e4.

15. Vigers, G., M. Coue, and J. McIntosh, Fluorescent microtubules break up under illumination. The Journal of cell biology, 1988. 107(3): p. 1011–1024.

16. Vemu, A., et al., Structure and Dynamics of Single-isoform Recombinant Neuronal Human Tubulin* Journal of Biological Chemistry, 2016. 291(25): p. 12907–12915.

17. Simmert, S., et al., LED-based interference-reflection microscopy combined with optical tweezers for quantitative three-dimensional microtubule imaging. Optics express, 2018. 26(11): p. 14499–14513.

18. Nong, D., et al., Integrated multi-wavelength microscope combining TIRFM and IRM modalities for imaging cellulases and other processive enzymes. Biomedical Optics Express, 2021. 12(6): p. 3253–3264.

19. Curtis, A., The mechanism of adhesion of cells to glass: a study by interference reflection microscopy. The Journal of cell biology, 1964. 20(2): p. 199–215.

20. Tuna, Y., A. Al-Hiyasat, and J. Howard, Imaging Dynamic Microtubules and Associated Proteins by Simultaneous Interference-Reflection and Total-Internal-Reflection-Fluorescence Microscopy. arXiv preprint arXiv:2201.07911, 2022.

21. Edelstein, A.D., et al., Advanced methods of microscope control using μManager software. Journal of biological methods, 2014. 1(2).

22. Ziółkowska, N.E. and A. Roll-Mecak, In vitro microtubule severing assays, in Adhesion Protein Protocols. 2013, Springer. p. 323-334.

23. Saper, G. and H. Hess, Kinesin-propelled label-free microtubules imaged with interference reflection microscopy. New Journal of Physics, 2020. 22(9): p. 095002.

24. Luchniak, A., et al., Dynamic microtubules slow down during their shrinkage phase. Biophysical Journal, 2023. 122(4): p. 616–623.

25. Ciorîță, A., et al., Single depolymerizing and transport kinesins stabilize microtubule ends. Cytoskeleton, 2021. 78(5): p. 177–184.

26. Jain, K., et al., Polymerization kinetics of tubulin from mung seedlings modeled as a competition between nucleation and GTP-hydrolysis rates. Cytoskeleton, 2021. 78(9): p. 436–447.

27. McCormick, L.A., et al., Interface-acting nucleotide controls polymerization dynamics at microtubule plus-and minus-ends. Elife, 2024. 12: p. RP89231.

28. Molines, A.T., et al., Physical properties of the cytoplasm modulate the rates of microtubule polymerization and depolymerization. Developmental Cell, 2022. 57(4):p. 466–479. e6.

